# Modeling competitive transplantation using HLA-mismatched human hematopoietic stem cells

**DOI:** 10.64898/2026.03.18.712629

**Authors:** Abdullahi Mobolaji Idowu, James Ropa, Stephanie Hurwitz

**Affiliations:** Department of Pathology and Laboratory Medicine, Indiana University School of Medicine, Indianapolis, IN, USA; Melvin and Bren Simon Comprehensive Cancer Center, Indianapolis, IN, USA; Department of Medical and Molecular Genetics, Indiana University School of Medicine, Indianapolis, IN, USA

**Keywords:** Hematopoietic stem cells, competitive transplantation, repopulating unit, HLA, bone marrow, chimerism

## Abstract

**Background:** Competitive transplantation is essential for defining intrinsic repopulating capacity of murine hematopoietic stem and progenitor cells (HSPCs), yet comparable assays for human cells have been limited by the lack of a robust *in vivo* platform.

**Methods:** Here, we describe a novel competitive transplantation method in humanized NOD.Cg-*Kit*^*W-41J*^ *Tyr* ^+^ *Prkdc*^*scid*^ *Il2rg*^*tm1Wjl*^/ThomJ (NBSGW) mice that enables simultaneous engraftment and longitudinal tracking of distinct human grafts within a shared microenvironment.

**Results:** Using human leukocyte antigen-mismatched donor CD34+ cells, this method facilitates standard flow cytometry panels to track multiple donor cell chimerism, lineage output, and HSPC composition. The experimental framework may be adapted to different mouse models, conditioning strategies, donor sources, and treatments.

**Conclusions:** Overall, this humanized competitive repopulation assay fills a critical translational gap and offers a flexible foundation for advancing mechanistic discovery in human hematopoietic biology and improving clinical strategies for stem cell transplantation.

## Background

Hematopoietic stem and progenitor cell (HSPC) transplantation is a therapeutic strategy to treat a number of hematologic malignancies, bone marrow (BM) failure syndromes, and congenital immunodeficiencies. Despite growing access, the procedure is associated with high rates of morbidity and mortality (1). Modeling transplantation in mice provides a powerful translational framework for improving outcomes in human HSPC transplantation. In particular, competitive transplantation, where two populations of donor cells are transplanted into the same recipient mouse, is a key functional assay to compare the intrinsic repopulating capacity of HSPCs in vivo. Pioneered in the 1990s, competitive repopulation assays remain the gold standard for evaluating and quantitating intrinsic properties of HSPCs that promote durable engraftment and long-term hematopoietic recovery (2). Many clinically relevant concepts, including graft potency (3,4), stem cell aging (5,6), competitive disadvantage after inflammatory or genotoxic stress (7–9), and the impact of co-morbidities such as donor obesity on HSPC performance (10), were first uncovered through competitive repopulation studies.

Murine modeling of competitive transplantation has broadly relied on use of congenic mouse strains (C57Bl6 background: CD45.2^+^; B6.SJL-PtprcaPep3b/BoyJ: CD45.1^+^). However, translation to human models has largely been constrained by species-specific differences and the absence of a robust system for simultaneous *in vivo* comparison of human HSPC populations. A few studies have attempted to model human competitive transplantation through implanted fetal bones (11), lentiviral labeling with fluorescent reporters (12), and sex-mismatched cells (13). Notably, these studies have predominantly relied on the use of cord blood (CB)-derived HSPCs.

Here we detail a protocol for performing functional competitive transplantation assays in NOD.Cg-*Kit*^*W-41J*^ *Tyr* ^+^ *Prkdc*^*scid*^ *Il2rg*^*tm1Wjl*^/ThomJ (NBSGW) mice using HLA-mismatched human CD34^+^ cells derived from BM, offering robust bone marrow and splenic engraftment to quantitate repopulation capacity.

## Methods

### Animals

NOD.Cg-KitW-41J Tyr + Prkdcscid Il2rgtm1Wjl/ThomJ (NBSGW; Jackson strain #026622) female mice were purchased from Jackson Laboratories and housed in a pathogen-free facility with ad libitum access to food and water.

#### HSPC culture

HLA-mismatched (HLA-A*02 or HLA-B*08) human BM-derived CD34+ cells were purchased from Ossium Health. Cells were thawed at 37°C in a water bath and washed by resuspending in serum free StemSpan™ SFEM II media (STEMCELL Technologies), followed by centrifuging twice at 400 × g for 10 min at 4 °C. Cells were grown in *ex vivo* culture for 4 days in complete medium composed of StemSpan™ SFEM II (STEMCELL Technologies) supplemented with 100 IU penicillin-streptomycin (Gibco), 100 ng/mL IL-6 (Peprotech), 100 ng/mL Flt3 ligand (PeproTech), 100 ng/mL SCF (PeproTech), 100 ng/mL TPO (PeproTech), 1 μM StemRegenin1 (STEMCELL Technologies), and 35 nM UM171 (APExBIO Technology) (14).

#### Bone marrow transplantation

Twenty-four hours prior to transplantation, 8-week-old NBSGW mice were conditioned with 15 mg/kg busulfan. Equal numbers of HLA-A*02 and HLA-B*08 were pooled for a total of 2×10^5 cells, and retro-orbitally injected in a final volume of 200uL sterile PBS. Peripheral blood (PB) sampling and BM aspiration were used to monitor human chimerism after transplantation. At 12-weeks post-transplantation, mice were euthanized in a CO_2_ chamber, followed by a cervical dislocation, and surgical removal of the spleen and bilaterial femurs and tibias. Single-cell suspensions of BM cells were collected by flushing the tibias and femurs with a 3mL syringe attached to a 26-gauge needle using Hank’s Balanced Salt Solution (HBSS) (Gibco). Tissue clumps were removed by passing the solution several times through an 18-guage needle, then filtering through a 70μm mesh strainer. Cells were centrifuged at 400 x g for 5 minutes before resuspending in 1mL 1X red blood cell (RBC) Lysis buffer (Biolegend) and incubating at 4°C for 10 minutes. Cells were washed with PBS by centrifugation at 400 x g for 5 minutes prior to antibody staining. Spleens were placed in a tissue culture plate with cold PBS and mechanically dissociated using the plunger end of a 3-mL syringe. The resulting cell suspension was passed through a 70µm cell strainer, before RBC lysis and washing as described above.

#### HSPC immunophenotyping

BM and spleen cells were stained with a panel of antibodies to examine chimerism and lineage composition; BM cells were additionally stained with antibodies to analyze HSPC subpopulations. An example of recommended antibody clones and concentrations are listed in **Tables 1-2**. Cells were incubated for 30 minutes at 4 °C with antibodies in 100μL PBS in 5mL FACS tubes, before washing in PBS by 400 x g for 5 minutes, and resuspension in PBS for flow cytometry. Flow cytometric analysis was performed using a FACSAria Fusion Cell Sorter or BD FACSCanto II flow cytometer.

**Table 1.** Antibody panel for human lineage analysis.

| Cell surface marker | Fluorophore | Clone | Manufacturer | Volume (μL) per 100μL in PBS |
| --- | --- | --- | --- | --- |
| Viability | FVS510 |  | BD Biosciences | 0.1 |
| mCD45 | PE-Cy7 | I3/2.3 | Biolegend | 2 |
| hCD45 | PE | HI30 | Biolegend | 2 |
| HLA-A02 | APC-Fire750 | BB7.2 | Biolegend | 2 |
| HLA-B08 | APC | REA145 | Miltenyi Biotec | 2 |
| CD3 | BV421 | SK7 | BD Biosciences | 1 |
| CD33 | BB515 | WM53 | BD Biosciences | 1 |
| CD19 | BB700 | SJ25C1 | BD Biosciences | 1 |

**Table 2.** Antibody panel for human HSPC subpopulation analysis.

| Cell surface marker | Fluorophore | Clone | Manufacturer | Volume (μL) per 100μL in PBS |
| --- | --- | --- | --- | --- |
| HLA-A02 | APC-Fire750 | BB7.2 | Biolegend | 2 |
| HLA-B08 | APC | REA145 | Miltenyi Biotec | 2 |
| CD34 | PE | 581 | Biolegend | 1 |
| CD38 | BV711 | HIT2 | Biolegend | 1 |
| CD45RA | FITC | HI100 | Biolegend | 1 |
| CD49f | BV421 | GoH3 | Biolegend | 1 |

#### Statistical analyses

Group comparisons for continuous variables were performed using t-tests for two groups. A p-value of <0.05 was considered significant. All statistical testing was performed in GraphPad Prism 10.6.0, R 4.5.0, or Microsoft Excel.

## Results

Flow cytometric panels to distinguish HLA-specific donor chimerism, lineage output, and HSPC subpopulation composition were validated using commercially available antibodies (**Table 1-2**). These panels allowed for distinguishment of mouse versus human cell chimerism and human HLA-specific donor chimerism (**Figure 1A-B**). We also assessed HLA-specific output of B-cells, myeloid cells, and T-cells in the spleen and BM, as well as CD34^+^ CD38^+^ CD45RA^+/-^ CD49f^+/-^ precursors in the BM (**Figure 1C-G**).

**Figure 1.**
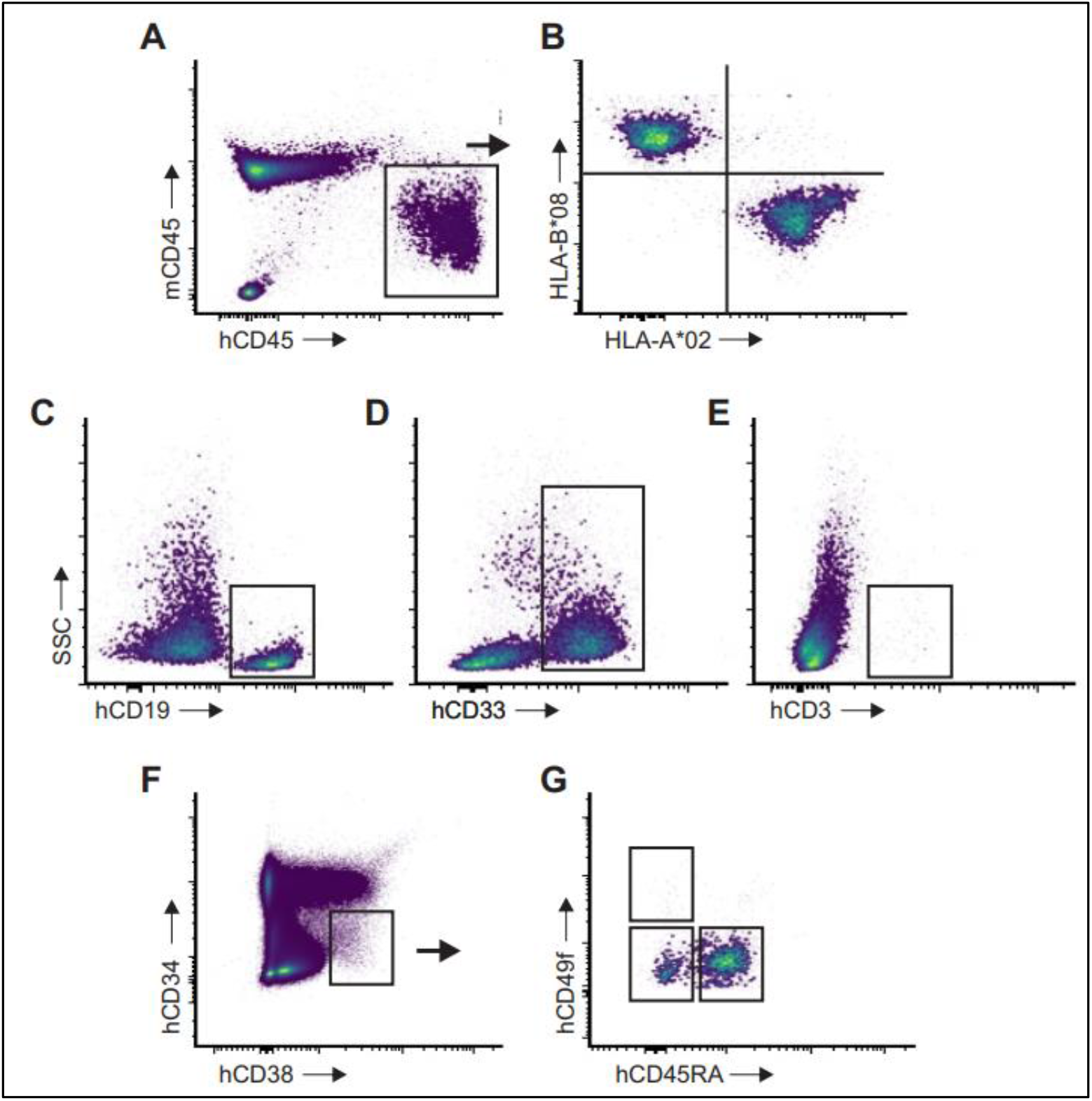
Flow cytometry gating strategies for tracking donor-specific repopulation. A) Representative plot showing bone marrow (BM) human (h) versus mouse (m) CD45 chimerism. B) HLA-specific antibodies were used to fluorescently tag donor cell populations after gating hCD45 cells. Reconstitution of BM (C) B-cells, (D) myeloid cells, and (E) T-cells within HLA-typed donors. F) Gating of CD34^+^ CD38^-^ precursors in engrafted BMs. G) Delineation of BM CD45RA^+/-^ CD49f^+/-^ stem and progenitor cells.

To account for intrinsic differences in repopulation capacity between HLA-mismatched donors, an experimental schematic is represented in **Figure 2A**. An example of how treatment effects may be calculated across donor cells with unequal baseline engraftment capacities is shown in **Table S1**. In this example, measured (M) versus expected (E) chimerism values can be calculated based on baseline donor differences to compute an estimated treatment effect. Appropriate statistical comparison testing may be performed on (M-E) compared to (E-E) values for each donor (15).

**Figure 2.**
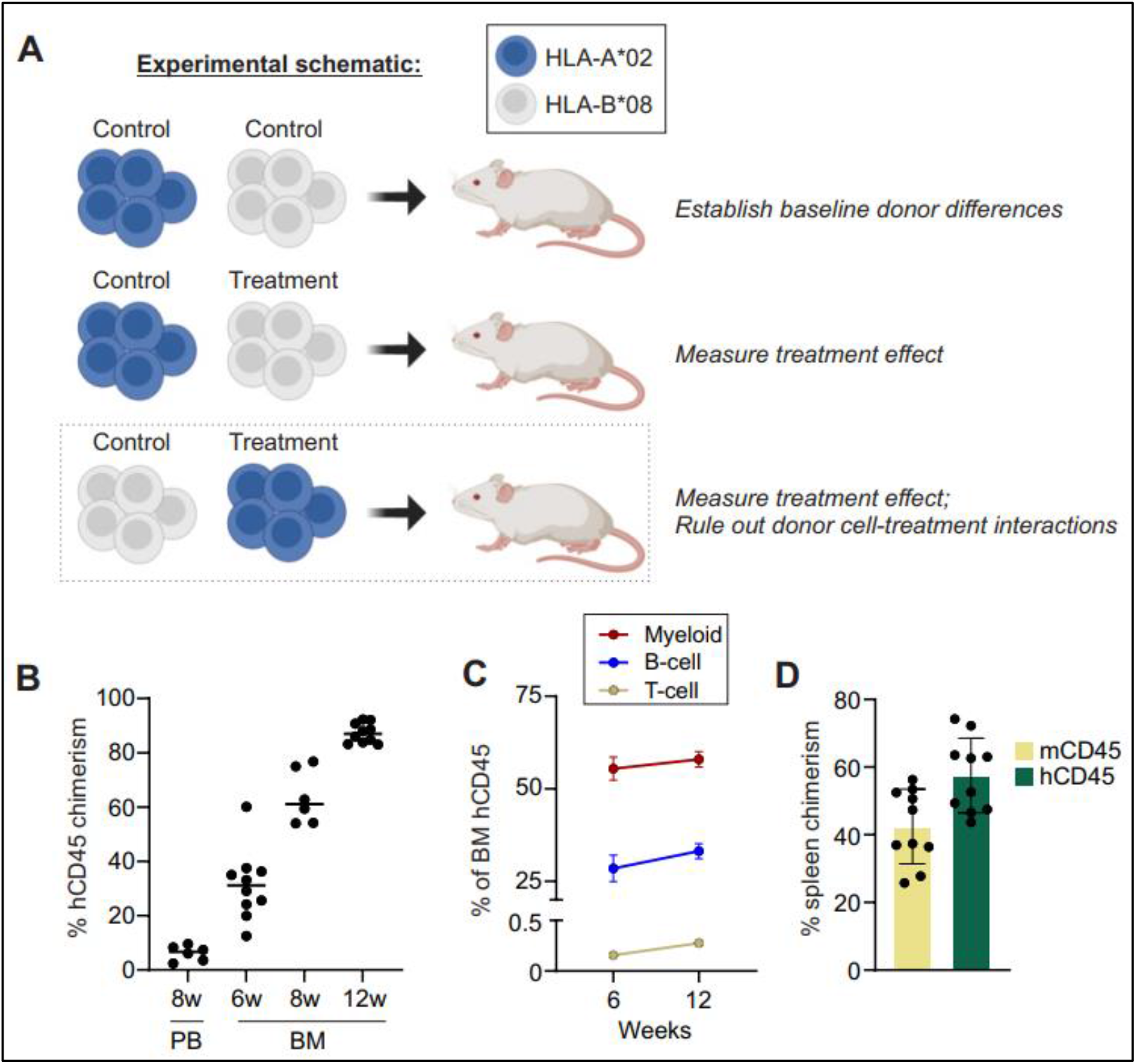
Experimental schematic for double graft xenotransplantation. A) Example schematic of experimental design to conduct HLA-mismatched competitive transplantations. B) Total percent of peripheral blood (PB) or bone marrow (BM) human (h) CD45^+^ cells, including both HLA types across weeks (w). C) Total percent of hCD45^+^ cells comprised of respective mature cells in engrafted BM. D) Relative mouse and human splenic chimerism at 12 weeks post-transplant.

In this study, we utilized BM CD34^+^ cells from HLA-typed donors: HLA-A*02 and HLA-B*08. As many researchers rely on short-term culture of cells for drug treatment or gene therapy, donor cells were cultured in cytokine-rich medium for four days before transplantation. For comparison of donor cell repopulation, humanized NBSGW mice were tracked for 12 weeks. In line with prior studies, levels of human cells in the PB of NBSGW mice were low (<10%) across groups at 8 weeks (**Figure 2B**) (9). Therefore, BM aspiration was performed at 6 and 8 weeks post-transplant, before endpoint analysis of the BM and spleen at 12 weeks. Total human BM chimerism steadily inclined between 6 and 12 weeks, with an average of 87% at week 12 (**Figure 2B**). Lineage output at both timepoints was myeloid-predominant, with very low levels of T-cell repopulation (**Figure 2C**). At 12 weeks, human chimerism in the spleen averaged 58% (**Figure 2D**).

HLA-A*02 and HLA-B*08 BM chimerism differed significantly at both 6 and 12 weeks in the BM and spleen, supporting the need for careful experimental design (**Figure 3A-B**). With exception of T-cell repopulation, lineage composition did not significantly differ between HLA-mismatched donors (**Figure 3C**). Similar to total donor cell chimerism, relative proportions of early progenitor cell frequencies were notably higher in the HLA-A*02 donor fraction (**Figure 3D-E**).

**Figure 3.**
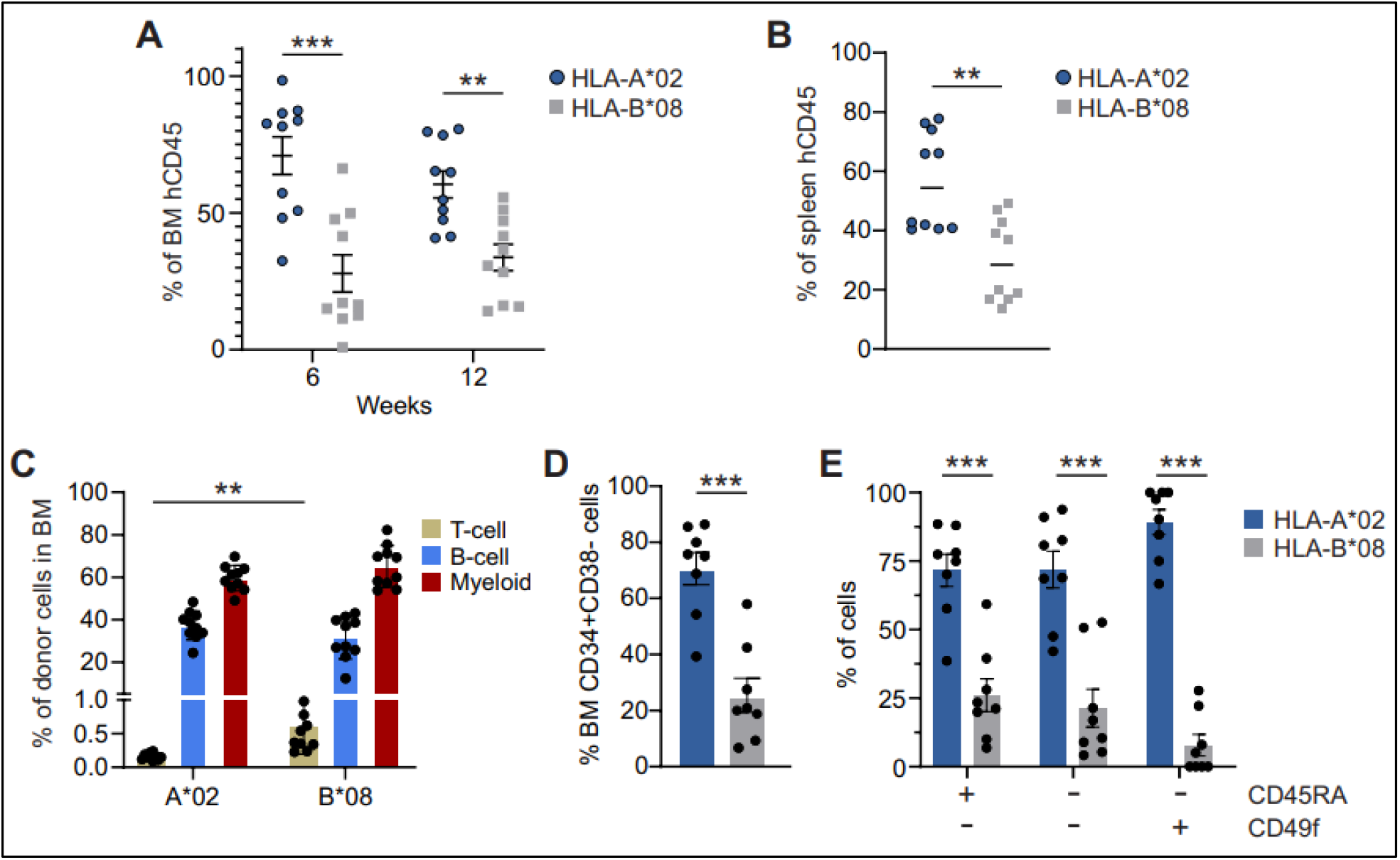
HLA-mismatched competitive transplantation allows dual donor cell tracking. HLA-specific (A) BM and (B) spleen chimerism, out of total human (h) CD45^+^ cells. C) Frequency of BM myeloid and lymphoid cells within each donor graft. HLA-typed donor contribution to BM (D) CD34^+^ CD38^-^ and (E) CD45RA^+/-^ CD49f^+/-^ stem and progenitor cells.

## Discussion

Here we describe a novel technique for assessing competitive hematopoietic engraftment in a humanized xenograft mouse model, addressing key limitations of existing transplantation assays and providing new opportunities to interrogate human HSPC biology in vivo. Use of HLA-mismatched donor cells facilitates a simple flow cytometry-based quantitative measure to capture comparative engraftment kinetics and graft output that may arise from genetic perturbation, drug exposure, or inherent donor differences. Further cell sorting may be performed to allow for downstream assays. Several HLA-targeting, oligonucleotide-conjugated antibodies are commercially available, and could also enable multiomic studies of engrafted HSPCs without cell sorting. We have previously demonstrated the utility of these competitive transplantations to examine the impact of *ex vivo* manipulation on intrinsic HSPC function (8).

HLA matching is a key selection criterion for HSPC donors, and is a strong predictive factor for transplantation success (10). While the impact of HLA-mismatch has been well studied, little is known about HLA-driven variability in the absence of donor-recipient mismatch. Given the repopulation differences seen between HLA types in our study, this method may be adopted to more broadly evaluate the impact of donor HLA type on intrinsic graft potency and reconstitution capacity. In addition, these studies may be helpful in elucidating mechanisms of xenogeneic graft-versus-host-disease (GVHD) and identifying which HLA types may be more prone to contribute to both GVHD and graft-versus-tumor effects.

Competitive transplantation modeling using fetal HSPCs is particularly relevant, given that double-unit CB transplantation is clinically used for patients without a better donor option. However, in the majority of cases where two separate CB units are administered as a graft, only one unit persists in recipients as a long-term source of hematopoiesis (11). Given that many researchers are already utilizing CB as a graft source in noncompetitive xenograft transplantations, our proposed technique may facilitate future mechanistic investigation into non-immunologic causes of single graft dominance.

In this study, we utilized the NBSGW mouse strain, as it has been previously demonstrated by our lab and others to facilitate high levels of human donor cell engraftment without myeloablative conditioning or irradiation (8,12). A high degree of human cell chimerism is optimal to sufficiently compare two or more donors, particularly when the resolution of rare progenitor cell frequencies is desired. In addition,

NBSGW mice support improved and physiologically relevant myeloid repopulation compared to other immunocompromised strains (9,12). We note that in our hands, human PB chimerism was substantially lower than frequencies of human hematopoietic cells in the BM of transplanted NBSGW mice. Others have reported variable levels of PB chimerism in these mice using different conditioning strategies, cell doses, and fetal HSPCs (9,13,14). Optimizing conditions may allow researchers to use PB sampling to track longitudinal chimerism in NBSGW mice and avoid more technically challenging BM aspirates. Comparison of NOD.Cg-Prkdc^scid^ Il2rg^tm1Wjl^/SzJ (NSG), NSG-SGM3 (NSGS), MISTRG, and NBSGW mice have been reported previously, each with advantages and disadvantages that may be considered (9,15). Overall, we believe tracking HLA-mismatched donor cells will facilitate competitive transplantation studies in researchers’ strain of choice for their designed experiments and lead to broader application.

In addition, future studies testing alternative donor cell sources may be warranted. Here, we establish a baseline protocol for the use of short-term ex vivo cultured BM CD34+ cells; however, this could be adopted for the use of fresh cells, mobilized PB, or CB HSPCs. HLA typing of donor cells by researchers, including by serology, polymerase chain reaction, Sanger sequencing, or next-generation sequencing, may be performed using commercially available assays. While we demonstrate the feasibility of tracking HLA-A*02 and HLA-B*08 serotypes, other available HLA-specific antibodies can facilitate the use of alternative serotypes. Finally, we offer a limited panel of antibodies tested as proof-of-concept to examine HSPC subpopulations; however, this panel may be expanded to analyze other discrete subpopulations.

## Conclusions

Taken together, we offer a flexible proof-of-concept technique that allows for future optimization of mouse strain, cell source, HLA type, and treatments to dissect human HSPC fitness and donor-associated variability within a controlled *in vivo* environment. As humanized models continue to evolve, this platform can serve as a foundational technique to allow *in vivo* graft tracking for mechanistic discovery and translational application.

## List of abbreviations

BM: bone marrow
CB: Cord blood
HBSS: Hank’s Balanced Salt Solution
HLA: Human Leukocyte Antigen
HSPC: hematopoietic stem and progenitor cell
PB: peripheral blood
PBS: phosphate-buffered saline

## Declarations

### Ethics approval and consent to participate

Animal experiments were approved by the Institutional Animal Care and Use Committee (IACUC) and conducted in accordance with the National Institutes of Health’s Guide for the Care and Use of Laboratory Animals. Human HSPCs were obtained from deidentified products purchased from Ossium Health. Patient consent was not obtained at the time of our purchase.

### Consent for publication

Not applicable.

### Availability of data and materials

The datasets generated and analyzed during the current study are available from the corresponding author on reasonable request.

### Authors’ contributions

S.N.H. conceived of and supervised the study, and collected data. A.M.I., J.R., and S.N.H. interpreted data. All authors wrote, reviewed, and edited the final manuscript.

## Acknowledgements

Thanks to Aidan Hurwitz for assistance in statistical design of experiments and Seul Jung for experimental coordination.

## References

1. Bolon Y, Atshan R, Allbee-Johnson M, Estrada-Merly N, Lee S. Current use and outcome of hematopoietic stem cell transplantation: CIBMTR summary slides, 2022.

2. Szilvassy SJ, Humphries RK, Lansdorp PM, Eaves AC, Eaves CJ. Quantitative assay for totipotent reconstituting hematopoietic stem cells by a competitive repopulation strategy. Proc Natl Acad Sci. 1990 Nov;87(22):8736–40.

3. Morrison SJ, Weissman IL. The long-term repopulating subset of hematopoietic stem cells is deterministic and isolatable by phenotype. Immunity. 1994 Nov 1;1(8):661–73.

4. Ema H, Sudo K, Seita J, Matsubara A, Morita Y, Osawa M, et al., Quantification of self-renewal capacity in single hematopoietic stem cells from normal and Lnk-deficient mice. Dev Cell. 2005 Jun;8(6):907–14.

5. Rossi DJ, Bryder D, Zahn JM, Ahlenius H, Sonu R, Wagers AJ, et al., Cell intrinsic alterations underlie hematopoietic stem cell aging. Proc Natl Acad Sci U S A. 2005 Jun 28;102(26):9194–9.

6. Chambers SM, Shaw CA, Gatza C, Fisk CJ, Donehower LA, Goodell MA. Aging hematopoietic stem cells decline in function and exhibit epigenetic dysregulation. PLoS Biol. 2007 Aug;5(8):e201.

7. Baldridge MT, King KY, Goodell MA. Inflammatory signals regulate hematopoietic stem cells. Trends Immunol. 2011 Feb;32(2):57–65.

8. Essers MAG, Offner S, Blanco-Bose WE, Waibler Z, Kalinke U, Duchosal MA, et al., IFNα activates dormant haematopoietic stem cells in vivo. Nature. 2009 Apr;458(7240):904–8.

9. Walter D, Lier A, Geiselhart A, Thalheimer FB, Huntscha S, Sobotta MC, et al., Exit from dormancy provokes DNA-damage-induced attrition in haematopoietic stem cells. Nature. 2015 Apr;520(7548):549–52.

10. Lee JM, Govindarajah V, Goddard B, Hinge A, Muench DE, Filippi MD, et al., Obesity alters the long-term fitness of the hematopoietic stem cell compartment through modulation of Gfi1 expression. J Exp Med. 2018 Feb 5;215(2):627–44.

11. Rosler ES, Brandt JE, Chute J, Hoffman R. An in vivo competitive repopulation assay for various sources of human hematopoietic stem cells. Blood. 2000 Nov 15;96(10):3414–21.

12. Yahata T, Ando K, Miyatake H, Uno T, Sato T, Ito M, et al., Competitive Repopulation Assay of Two Gene-Marked Cord Blood Units in NOD/SCID/γcnull Mice. Mol Ther. 2004 Nov 1;10(5):882–91.

13. Tatekawa T, Ogawa H, Kawakami M, Oka Y, Yasukawa K, Sugiyama H, et al., A novel direct competitive repopulation assay for human hematopoietic stem cells using NOD/SCID mice. Cytotherapy. 2006 Jan 1;8(4):390–8.

14. Hurwitz SN, Jung SK, Kobulsky DR, Fazelinia H, Spruce LA, Baltasar-Pérez E, et al., Neutral sphingomyelinase blockade enhances hematopoietic stem cell fitness through an integrated stress response. Blood. 2023 Sep 12;142(20):1708–23.

15. Seiffert S, Weber S, Sack U, Keller T. Use of logit transformation within statistical analyses of experimental results obtained as proportions: example of method validation experiments and EQA in flow cytometry. Front Mol Biosci. 2024 Jul 11;11:1335174.

16. Sippel TR, Radtke S, Olsen TM, Kiem HP, Rongvaux A. Human hematopoietic stem cell maintenance and myeloid cell development in next-generation humanized mouse models. Blood Adv. 2019 Jan 29;3(3):268–74.

17. Choo S, Wolf CB, Mack HM, Egan MJ, Kiem HP, Radtke S. Choosing the right mouse model: comparison of humanized NSG and NBSGW mice for in vivo HSC gene therapy. Blood Adv. 2024 Feb 27;8(4):916–26.

18. Hess NJ, Lindner PN, Vazquez J, Grindel S, Hudson AW, Stanic AK, et al., Different Human Immune Lineage Compositions Are Generated in Non-Conditioned NBSGW Mice Depending on HSPC Source. Front Immunol [Internet]. 2020 Oct 19 [cited 2025 Dec 18];11. Available from: https://www.frontiersin.org/journals/immunology/articles/10.3389/fimmu.2020.573406/full

19. Petersdorf EW, Gooley TA, Anasetti C, Martin PJ, Smith AG, Mickelson EM, et al., Optimizing outcome after unrelated marrow transplantation by comprehensive matching of HLA class I and II alleles in the donor and recipient. Blood. 1998 Nov 15;92(10):3515–20.

20. Sasazuki T, Juji T, Morishima Y, Kinukawa N, Kashiwabara H, Inoko H, et al., Effect of matching of class I HLA alleles on clinical outcome after transplantation of hematopoietic stem cells from an unrelated donor. Japan Marrow Donor Program. N Engl J Med. 1998 Oct 22;339(17):1177–85.

21. Lee SJ, Klein J, Haagenson M, Baxter-Lowe LA, Confer DL, Eapen M, et al., High-resolution donor-recipient HLA matching contributes to the success of unrelated donor marrow transplantation. Blood. 2007 Dec 15;110(13):4576–83.

22. Ishiwata K, Ota H, Nishida A, Tsuji M, Yamamoto H, Yamamoto G, et al., Impact of HLA Haplotype Matching On Engraftment in Reduced Intensity Cord Blood Transplantation. Blood. 2012 Nov 16;120(21):3043.

23. Morishima Y, Kashiwase K, Matsuo K, Azuma F, Morishima S, Onizuka M, et al., Biological significance of HLA locus matching in unrelated donor bone marrow transplantation. Blood. 2015 Feb 12;125(7):1189–97.

24. Sasazuki T, Juji T, Morishima Y, Kinukawa N, Kashiwabara H, Inoko H, et al., Effect of matching of class I HLA alleles on clinical outcome after transplantation of hematopoietic stem cells from an unrelated donor. Japan Marrow Donor Program. N Engl J Med. 1998 Oct 22;339(17):1177–85.

25. Petersdorf EW, Gooley TA, Anasetti C, Martin PJ, Smith AG, Mickelson EM, et al., Optimizing outcome after unrelated marrow transplantation by comprehensive matching of HLA class I and II alleles in the donor and recipient. Blood. 1998 Nov 15;92(10):3515–20.

26. Morishima Y, Sasazuki T, Inoko H, Juji T, Akaza T, Yamamoto K, et al., The clinical significance of human leukocyte antigen (HLA) allele compatibility in patients receiving a marrow transplant from serologically HLA-A, HLA-B, and HLA-DR matched unrelated donors. Blood. 2002 Jun 1;99(11):4200– 6.

27. Petersdorf EW, Malkki M, Hsu K, Bardy P, Cesbron A, Dickinson A, et al., 16th IHIW: international histocompatibility working group in hematopoietic cell transplantation. Int J Immunogenet. 2013 Feb;40(1):2–10.

28. Ehx G, Somja J, Warnatz HJ, Ritacco C, Hannon M, Delens L, et al., Xenogeneic Graft-Versus-Host Disease in Humanized NSG and NSG-HLA-A2/HHD Mice. Front Immunol. 2018;9:1943.

29. Barker JN, Weisdorf DJ, DeFor TE, Blazar BR, McGlave PB, Miller JS, et al., Transplantation of 2 partially HLA-matched umbilical cord blood units to enhance engraftment in adults with hematologic malignancy. Blood. 2005 Feb 1;105(3):1343–7.

30. McIntosh BE, Brown ME, Duffin BM, Maufort JP, Vereide DT, Slukvin II, et al., Nonirradiated NOD,B6.SCID Il2rγ™/™KitW41/W41 (NBSGW) Mice Support Multilineage Engraftment of Human Hematopoietic Cells. Stem Cell Rep. 2015 Jan 15;4(2):171–80.

31. Leonard A, Yapundich M, Nassehi T, Gamer J, Drysdale CM, Haro-Mora JJ, et al., Low-Dose Busulfan Reduces Human CD34+ Cell Doses Required for Engraftment in c-kit Mutant Immunodeficient Mice. Mol Ther Methods Clin Dev. 2019 Nov 11;15:430–7.

32. Shultz LD, Lyons BL, Burzenski LM, Gott B, Chen X, Chaleff S, et al., Human Lymphoid and Myeloid Cell Development in NOD/LtSz-scid IL2Rγnull Mice Engrafted with Mobilized Human Hemopoietic Stem Cells1,2. J Immunol. 2005 May 1;174(10):6477–89.

33. Chen J, Liao S, Xiao Z, Pan Q, Wang X, Shen K, et al., The development and improvement of immunodeficient mice and humanized immune system mouse models. Front Immunol [Internet]. 2022 Oct 19 [cited 2025 Nov 21];13. Available from: https://www.frontiersin.org/journals/immunology/articles/10.3389/fimmu.2022.1007579/full

